# Dynamic chromatin regulatory landscape of human CAR T cell exhaustion

**DOI:** 10.1101/2021.04.02.438240

**Authors:** David G Gennert, Rachel C Lynn, Jeff M Granja, Evan W Weber, Maxwell R Mumbach, Yang Zhao, Zhana Duren, Elena Sotillo, William J. Greenleaf, Wing H Wong, Ansuman T Satpathy, Crystal L Mackall, Howard Y Chang

**Affiliations:** Center for Personal Dynamic Regulomes, Stanford University, Stanford, CA 94305; Department of Genetics, Stanford University, Stanford, CA 94305; Center for Cancer Cell Therapy, Stanford Cancer Institute, Stanford University School of Medicine, Stanford, CA 94305; Department of Statistics, Stanford University, Stanford, CA 94305; Department of Applied Physics, Stanford University, Stanford, CA 94305; Chan Zuckerberg Biohub, San Francisco, CA 94158; Department of Pathology, Stanford University School of Medicine, Stanford, CA 94305; Department of Pediatrics, Stanford University School of Medicine, Stanford, CA; Department of Medicine, Stanford University School of Medicine, Stanford, CA; Howard Hughes Medical Institute, Stanford University, Stanford, CA 94305

## Abstract

Dysfunction in T cells limits the efficacy of cancer immunotherapy^1–6^. We profiled the epigenome, transcriptome, and enhancer connectome of exhaustion-prone GD2-targeting HA-28z chimeric antigen receptor (CAR) T cells and control CD19-targeting CAR T cells, which present less exhaustion-inducing tonic signaling, at multiple points during their *ex vivo* expansion^7^. We found widespread, dynamic changes in chromatin accessibility and 3D chromosome conformation preceding changes in gene expression, notably at loci proximal to exhaustion-associated genes such as *PDCD1, CTLA4*, and *HAVCR2*, and increased DNA motif access for AP-1 family transcription factors, which are known to promote exhaustion. Although T cell exhaustion has been studied in detail in mouse, we find that the regulatory networks of T cell exhaustion differ between the species and involve distinct loci of accessible chromatin and cis-regulated target genes in human CAR T cell exhaustion. Deletion of exhaustion-specific candidate enhancers of *PDCD1* suppress the expression of PD-1 in an *in vitro* model of T cell dysfunction and in HA-28z CAR T cells, suggesting enhancer editing as a path forward in improving cancer immunotherapy.

## INTRODUCTION

Chimeric antigen receptor (CAR) T cells have proven to be an effective cancer immunotherapy strategy against B cell malignancies^2,4,8,9^, but T cell exhaustion has been identified as a major roadblock in development of universally effective CAR T cells^10–13^. First noted in models of chronic viral infection, T cell exhaustion occurs in the context of persistent antigen stimulation and is associated with limited T cell proliferation, cytokine production, and cell killing in response to further stimulation. T cell exhaustion is now recognized as a distinct epigenetic state, where fully exhausted T cells exhibit chromatin changes and cannot be reversed by checkpoint blockade but can be remodeled by cessation of cell signaling^14–16^. Exhaustion in CAR T cells can arise via high tumor burdens or tonic CAR signaling during *ex vivo* expansion, which severely limits therapeutic efficacy^10,17^.

Much of the current understanding of the genetic regulation underpinning T cell function^18–20^, generally, and T cell exhaustion^21–27^, specifically, come from mouse models of the immune system and chronic infection, while genomic studies in dysfunctional human T cells are only very recently emerging^15,21–23,28–34^ While we previously identified the dysregulation of AP-1 transcription factors as the driver of exhaustion in CAR T cells^7^, the dynamics of the chromatin state during the progression of T cell exhaustion and the genetic regulatory program underlying the progression remain unclear. Here, we present the chromatin landscape of exhaustion-prone GD2-targeting HA-28z and non-exhausted CD19-targeting CD19-28z CAR T cells throughout the development of the exhaustion phenotype.

Here we identify and differentiate the gene regulatory networks involved in the early development and later maintenance of the exhausted T cell phenotype, across various T cell subtypes found in CAR T cell cultures. As the genetic networks underlying the T cell response to antigen and T cell exhaustion are largely independent^25^, we can dissect the uniquely exhaustion-associated network and uncover coregulated modules of accessible chromatin loci, transcription factors, and dysregulated genes. The genomic features we identify may be used to inform targeted genomic using CRISPR/Cas9 to restrict expression of exhaustion markers such that the *ex vivo* maturation of CAR T cells does not induce T cell exhaustion prior to transfusion.

## RESULTS

### HA-28z CAR T cells become exhausted during *in vitro* culture and exhibit a unique chromatin signature

We have previously reported that human HA-28z CAR T cells, targeting the GD2 surface marker, exhibit tonic signaling and lead to an exhausted T cell phenotype following 10 days of transduction, in contrast to the non-exhausted phenotype of human CD19-28z CAR T cells, targeting the CD19 surface marker^7,16^. To elucidate the dynamics of the exhaustion-associated chromatin state, we profiled the chromatin accessibility landscape and transcriptome of human CAR T cells transduced with either the exhaustion-inducing HA-28z CAR or the non-exhausting CD19-28z CAR (Figure 1a).

**Figure 1.**
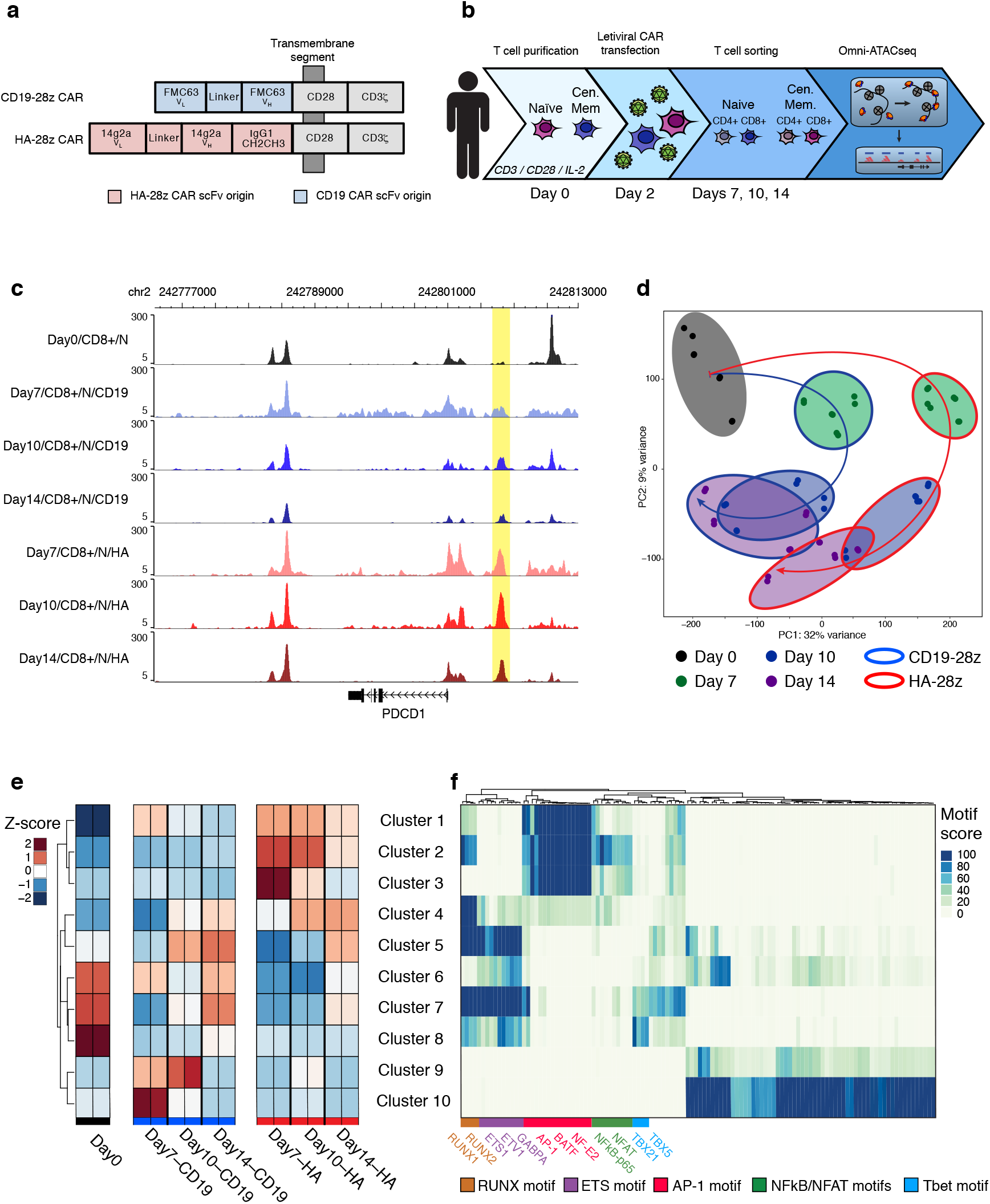
HA-28z CAR T cells exhibit a differentially accessible chromatin profile during the development of exhaustion. Donor T cells were collected and transfected with either the HA-28z or CDl9-28z CAR construct. (A) Schematic view of the two CAR constructs. (B) Experimental schematic outlining the time course of stimulation, transfection, and cell collection for ATAC-seq. (C) Differentially accessible chromatin at the PDCD1 locus in naïve CD8+ HA-28z and CD19-28z CAR T cells, shown as normalized ATAC-seq tracks. The element highlighted shows increased, sustained accessibility in HA-28z CAR T cells. (D) Principal component analysis (PCA) of all samples by global chromatin accessibility profile. PC1 (32% variance) separate HA-28z CAR T cells from CDl9-28z CAR T cells. (D) K-means clustering of top 5000 most variable chromatin accessibility peaks, colored by mean z-score within each cluster. (E) TF motif enrichment within each cluster from (D).

Throughout the 14-day *in vitro* maturation, the HA-28z CAR T cells more strongly expressed markers of exhaustion, including the inhibitory surface receptors PD-1, TIM-3, LAG3, CD39, CD45RA, CD62L, and CCR7; the secretion of activating cytokines IL-2 and IFNg; and decreased expansion, as compared to the CD19-28z CAR T cells (Supplemental Figure 1a-b). These phenotypic differences can be observed for some markers by Day 7, were most evident by day 10, and persist to Day 14. We collected HA-28z and CD19-28z CAR T cells starting on the day of purification and naïve/central memory sorting from a donor blood draw (Day 0) and subsequently on Days 7, 10, and 14 after CAR transduction on Day 2. On each collection day, cells were sorted into CD4+/naïve, CD4+/central memory, CD8+/naïve, and CD8+/central memory populations. RNA and chromatin were purified from each population, and RNA-seq and Omni-ATAC-seq were performed on each sample, respectively (Figure 1b, Supplemental Figure 1c-d).

Aggregating the ATAC-seq data reveals loci with differential chromatin accessibility both across time and across CAR type, including loci proximal to known T cell exhaustion marker gene *PDCD1* (Figure 1c). We found global differences in chromatin accessibility associated with the CAR type that exhibit dynamic genome-wide organization throughout the 14day maturation time course, with a notable contraction of differentially accessible loci in CD19-28z CAR T cells by the later time points, most prominently in the Days 10 and 14 CD19-28z CAR T cells (Supplemental Figure 2a-c). The chromatin profiles of the two CAR types exhibits the highest level of differential accessibility at Day 7 of the time course, before the differential increase in surface expression of exhaustion markers PD-1, CTLA4, and dysfunctional IL-2 secretion manifests. Although thousands of loci are differentially accessible throughout the time course, many are restricted to specific time points (Supplemental Figure 2d). Thus, the chromatin and gene regulatory changes precede the manifestation of the exhaustion phenotype, suggesting a causal role.

Clustering all ATAC-seq profiles reveals that CD4+, CD8+, naïve-derived, and central memory-derived HA-28z CAR T cells share high correlation among their global chromatin profiles across all time points (Supplemental Figure 2e). The Day 0 (prior to CAR transduction and activation) T cells form a cluster most closely associated with the late (Days 10 and 14) naïve CD19-28z CAR T cells. Principal component analysis (PCA) of global Omni-ATAC-seq profiles implicate the CAR type as the primary driver of differences in chromatin accessibility, as the PC1 axis separates the profiles of CAR T cells sharing a time point but differing in CAR type (Figure 1d). PCA suggests a cyclical pattern of global chromatin response to the *in vitro* maturation in CD19-28z CAR T cells, as their chromatin profiles approach the non-transduced Day 0 populations by Days 10 and 14. This pattern appears disrupted in exhausted HA-28z CAR T cells as their PCA trajectory leaves them more distant from the Day 0 populations.

### Global Patterns of Chromatin Accessibility Highlight the Role of AP-1 in Exhaustion

Performing hierarchical clustering on the 5000 ATAC-seq peaks, or accessible loci, with the greatest variance across normalized samples resulted in clusters of peaks differentially accessible in CD19-28z or HA-28z CAR T cells, many of which appear proximal to genes associated with T cell exhaustion and function, such as *PDCD1, CTLA4, HAVCR2, TIGIT, LAG3*, and *TOX*^31,35,36^, while also revealing that differential clusters of peaks associated with CD4+/CD8+ and naïve/central memory phenotypes remain a small fraction of the driving differences between cell populations (Supplemental Figure 2f).

Aggregating the accessibility signal within each cluster, we find 10 clusters of peaks with unique temporal dynamics of enrichment within the T cells across the time course (Figure 2e). Transcription factor motif analysis on the peaks within each cluster showed high enrichment in the CAR type-associated clusters for certain families of transcription factor motifs. Notably, the AP-1 motif is enriched in HA-28z CAR T cell peaks starting at Day 7 (clusters 1-3), suggesting a role for members of this TF family in the establishment of exhaustion, in accordance with our previous findings that inhibitory AP-1 family transcription factors promote exhaustion in human CAR T cells^7^, while RUNX and ETS family motifs are enriched in peaks that appear starting at Day 10 or 14 in HA-28z CAR T cells (clusters 4-5), suggesting a role in maintenance of exhaustion (Figure 2f).

**Figure 2.**
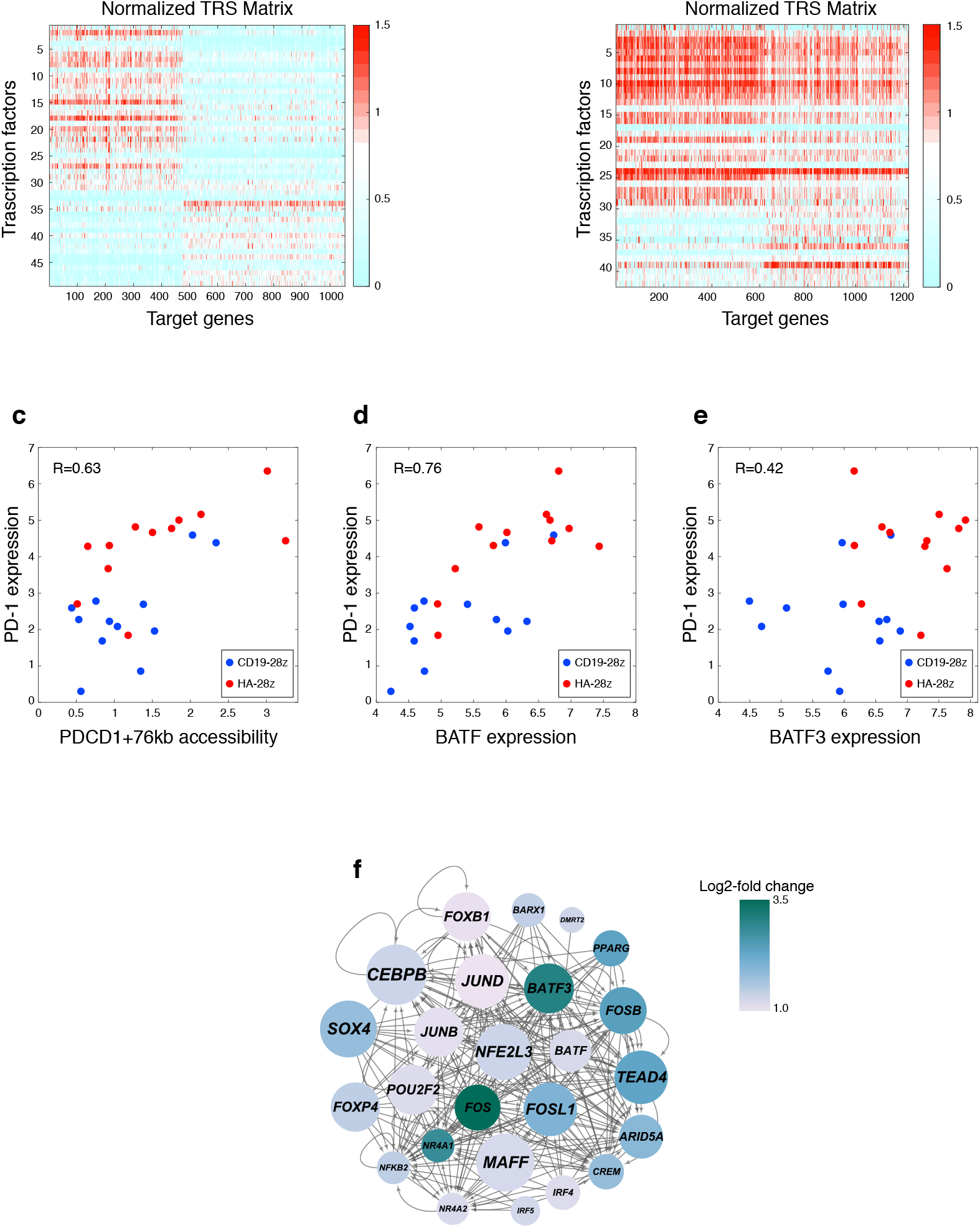
Integration of RNA-seq and ATAC-seq reveals independent regulatory gene-TF modules. (A) Representative transcription factor by gene regulatory score matrix as calculated by PECA2 for the Day 7/CD4+/Central Memory/HA-28z CAR T cells and (B) matching Day 7/CD4+/Central Memory/CD19-28z CAR T cells. (C) Correlation between the mRNA expression of PD-l and the normalized chromatin accessibility at one locus 76 kb upstream of the *PDCD1* transcription start site, identified by PECA2 as regulatory to PD-1. (D) The correlation between the mRNA expression of *PD-1* and mRNA expression of the transcription factors *BATF* and (E) *BATF3*. (F) PECA2 inferred gene regulatory network for the Day 7/CD4+/CM CAR T cells, showing TFs whose log2 fold-change between HA-28z and CD19-28z CAR T cells is >1. Node color presents log2 fold-change and node size represents enrichment score, or the geometric mean of fold-change and −log10(p-value)) on HA-specific genes and HA-specific regulatory elements.

### Integration of Transcriptome and Epigenome Identify Coregulated Regulatory Element-Gene Modules

We used the PECA2 and EnrichTF methods^37,38^ to integrate our multi-omic data set, incorporating RNA expression, chromatin accessibility, and time to identify coregulated modules of genetic regulatory elements, transcription factors, and target genes. Across the CAR T cells collected at Day 7 of *ex vivo* culture, 6501 DNA elements were identified as differentially accessible in HA-28z CAR T cells, and 698 genes were identified as differentially expressed in HA-28z CAR T cells over CD19-28z CAR T cells, both meeting the criteria of p<0.05 and foldchange of ≥1.2. A number of AP-1 family TFs are represented in the list of the 20 TFs with the highest Day 7 HA-28z-specific regulatory scores as called by EnrichTF, with NFE2L1, a bZIP TF known to interact with the AP1 complex at gene promoters^39,40^, and its close homolog NFE2L3^41^, as the highest-scoring TFs (Supplemental Table 1).

Across Day 10 CAR T cells, we found 4054 DNA elements and 458 target genes to be differentially accessible or expressed, respectively, in HA-28z CAR T cells, as compared to CD19-28z CAR T cells. Similarly to Day 7 CAR T cells, NFE2L1 and NFE2L3 were the highest-scoring HA-28z-specific regulatory TFs at Day 10, with other AP-1 TFs in the highest-scoring list as well (Supplemental Table 2).

We compared the TFs called by EnrichTF as differentially regulatory in HA-28z CAR T cells across time points to identify the TFs that play roles in gene regulation specifically at an early time point (Day 7) or later time point (Day 10) during the development of exhaustion in HA-28z CAR T cells. TFs such as NFE2L1, FOXP4, and JUND were found to be both significant regulators in the HA-28z CAR T cells and higher-scoring in the Day 7 TF analysis compared to the Day 10 TF analysis (Supplementary Table 3). Conversely, EGR3, BATF3, and ARID3B were among the TFs found to be significant regulators in the HA-28z CAR T cells and higher scoring in the Day 10 TF analysis compared to the Day 7 TF analysis (Supplementary Table 3).

The association of transcription factors to their target genes allows us to cluster these associations into regulatory modules by the strength of their associations in each sample using PECA2. The HA-28z CAR T cells, across CD4+/CD8+ and naïve/central memory subtypes, show two distinct and mutually exclusive regulator-gene modules in their significantly coregulated TF/target gene pairs (Figure 2a, Supplemental Figure 3a). The CD19-28z CAR T cells, however, do not show a strong breakdown of their TF/target gene associations into distinct modules (Figure 2b, Supplemental Figure 3b). Known exhaustion marker genes are found across both regulatory modules in HA-28z CAR T cells, including PDCD1 (Module 1) and CTLA4, LAG3, JUN, and JUNB (Module 2) (Supplemental Table 4). Transcription factors in the two regulatory modules include AP-1 TFs JUN, JUNB, and ATF3 (Module 1), BATF and BATF3 (Module 2) (Supplementary Table 5). NFE2L1 was found to be the highest-scoring TF at the Day 7 time point in Module 2, again suggesting its importance in the regulation of the exhaustion genetic program.

As an example of PECA2 identifying regulatory element/TF/target gene associations, we found a locus approximately 76 kb upstream of the *PDCD1* promoter, which contained a BATF/BATF3 consensus motif sequence. *BATF* mRNA expression correlated with *PD-1* mRNA expression (Figure 2c), more closely than BATF3 does (Figure 2d), while the chromatin accessibility of this locus also correlates with the expression of PD-1 (Figure 2e). Together, this suggests a role for BATF regulating expression of PD-1 by binding at the PDCD1-76kb locus. We can infer a broader gene regulatory network through PECA2 for each CD19-28z/HA-28z matched pair, such as the network that maintain the differentially expressed genes between the exhausted and non-exhausted Day 7/CD4+/CM CAR T cells, which highlights a central role for BATF, BATF3, several AP-1 factors, and NFE2L3 (Figure 2f).

### Human vs. Mouse T Cell Exhaustion Exhibit Distinct Chromatin Landscapes

The noncoding genome is under far less stringent constraint over evolutionary time, and DNA regulatory elements can rewire to produce a conserved gene expression outcome even among related mammalian species^42–44^. We compared our chromatin accessibility profiles in human T cells to the previously published mouse exhaustion datasets. It has been previously reported that loci proximal to the mouse *Pdcd1* gene are differentially accessible under prolonged, chronic stimulation, many of which share sequence homology with loci proximal to *PDCD1* in our CD19-28z and HA-28z CAR T cells (Figure 3a)^21^. In addition, the genomic features underlying the ATAC-seq peak sequences are quite similar across species and T cell exhaustion state (Supplemental Figure 4a).

**Figure 3.**
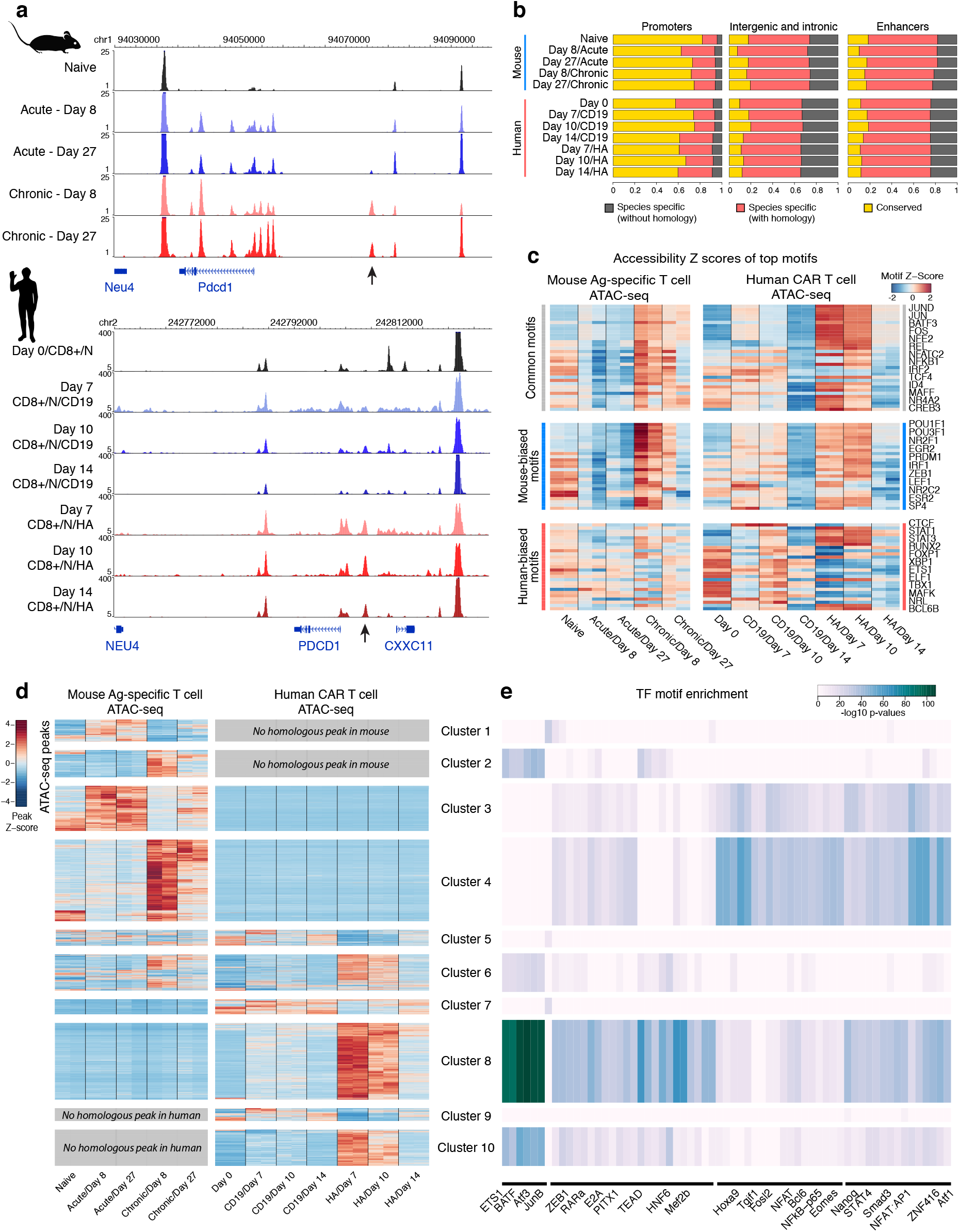
Exhausted human and mouse CAR T cells both share regulatory circuits and rely on species-restricted regulatory circuits. (A) Chromatin accessibility at the *PDCD1* locus in mouse- and human-derived CAR T cells under non-exhaustive (acute stimulation/CD19-28z CAR) and exhaustive (chronic stimulation/HA-28z CAR) conditions. Arrow highlights mouse and human elements that are accessible in the exhausted state. (B) Proportions of accessible chromatin loci determined to be accessible at conserved sequences between species (“Conserved”), accessible only in one species with sequence homology in the other species (“Species specific, mapped”), or accessible only in one species without sequence homology in the other species (“Species specific, unmapped”). (C) Enrichment of TF binding motifs across all the accessible chromatin loci in mouse- and human-derived CAR T cells, highlighting motifs that are common between species, have higher enrichment and variability in human, and have higher enrichment and variability in mouse. (D) Hierarchical clustering of normalized chromatin accessibility in individual loci across mouse- and human-derived CAR T cells, broken into clusters by the species in which the underlying sequence appears, and in which samples accessibility is enriched and differential between exhausted and non-exhausted cells. (E) The enrichment of TF binding motif sequences in each cluster of accessible loci.

However, the interspecies sequence conservation of the underlying genome varies with genomic feature. We found that the majority of peaks at gene promoters were found in both species, while the majority of peaks found at intergenic, intronic, and regions overlapping with the Fantom5 CD4+ T cell enhancer list^45^ were largely unique to either human or mouse, both with and without homologous sequences across species (Figure 3b).

To address whether the differences in chromatin profiles associated with T cell exhaustion between species is associated with specific transcription factors, we used transcription factor motif enrichment to identify the motifs enriched in peaks that are either shared across species, such as members of the AP-1 family, or are biased towards accessibility in only one of the species, such as the mouse-biased POU family and the human-biased STAT family (Figure 3c). We then performed differential analysis to identify exhaustion associated accessible chromatin loci across mouse and human samples by their normalized accessibility and identified clusters representing peaks differentially accessible in exhausted or non-exhausted T cells, with shared homology or not (Figure 3d). The majority of exhaustion-associated peaks were found to be species-specific– that is, most loci accessible in exhausted T cells were accessible in either (but not both) mouse or human T cells, most loci accessible in non-exhausted cells were similarly accessible in either (but not both) species, and few loci were accessible in both species in the corresponding cell types (Supplementary Figure 4b). A small fraction of peaks had reversed accessibility dynamics, with high accessibility in the exhausted T cells of one species and the non-exhausted T cells of the other species (Supplementary Figure 4b). As we observed in exhausted human CAR T cells (Supplementary Figure 2d), the thousands of loci associated with exhaustion in mouse T cells demonstrate an early, strong wave of increased accessibility followed by a gradual decrease in accessibility in a return to a more basal state (Figure 3d). Together, this analysis illustrates a high level of divergence in the regulatory programs underpinning T cell exhaustion in mouse and human through the utilization of largely unique enhancer sequences and, in some instances, evolving contrasting function for the same locus, while maintaining a characteristic chromatin signature and dynamic profile over time.

We performed transcription factor enrichment on each cluster of peaks, and notably observed AP-1 TF motifs most significantly enriched in peaks associated with the exhausted HA-28z human CAR T cells (Cluster 8) (Figure 3e). While several peak clusters represent loci accessible primarily in one species, the TF motifs enriched in those clusters are often shared between species. For example, Clusters 4 and 8 contain peaks accessible in exhausted mouse and human T cells, respectively, yet both have high enrichment for such TF motifs as RARa, STAT4, and SMAD3. Several TF motifs within these clusters do demonstrate species bias, however, such as high enrichment of HNF6 and MEF2B in the human exhaustion-biased cluster 8 and high enrichment of Hoxa9 and ZNF416 motifs in the mouse exhaustion-biased cluster 4. We also note that clusters composed of peaks without homologs in the other species have distinct TF motif enrichment patterns, suggesting significant higher-level divergence in the T cell exhaustion regulation program between mouse and human.

KEGG pathway enrichment analysis on GREAT annotations of exhaustion-associated clusters of peaks (clusters 2, 4, 6, 8, 10) reveal largely shared pathways for the accessible chromatin loci in exhausted T cells across species, including apoptosis, T cell receptor signaling, and JAK-STAT signaling, along with several mouse-biased pathways, such as the PD-L1/PD1 pathway, and the human-biased NFκB signaling pathway (Supplemental Figure 4c). Gene ontology analysis of the annotations of exhaustion-associated clusters indicates that the human and mouse shared exhaustion-related genomic loci are enriched for an immune cell activation and differentiation signal, while the human-specific exhaustion ATAC signature is notably enriched for regulation of cytokines IL-2 and IL-4 (Supplemental Figure 4d).

### HiChIP Identifies Distal Enhancers Associated With CAR T Cell Exhaustion

Many DNA regulatory elements are located tens to hundreds of kilobases away from its target genes, and 3D chromosome folding brings regulatory elements and target genes in spatial proximity while excluding intervening genes. To understand the 3D chromosome landscape, we performed HiChIP directed against histone H3 lysine 27 acetylation (H3K27ac), a histone mark associated with active enhancers, to construct the enhancer connectome^46^. We performed H3K27ac HiChIP and identified differential 3D chromatin looping in Day 0 starting T cell population, Day 10 exhausted CD8+/naïve HA-28z CAR T cells, and Day 10 non-exhausted CD8+/naïve CD19-28z CAR T cells. A number of gene promoters associated with T cell exhaustion lie within the top 5000 most variable 10 kb HiChIP loop anchor windows, including *PDCD1, HAVCR2, LAG3, TIGIT*, and *TOX* (Figure 4a).

**Figure 4.**
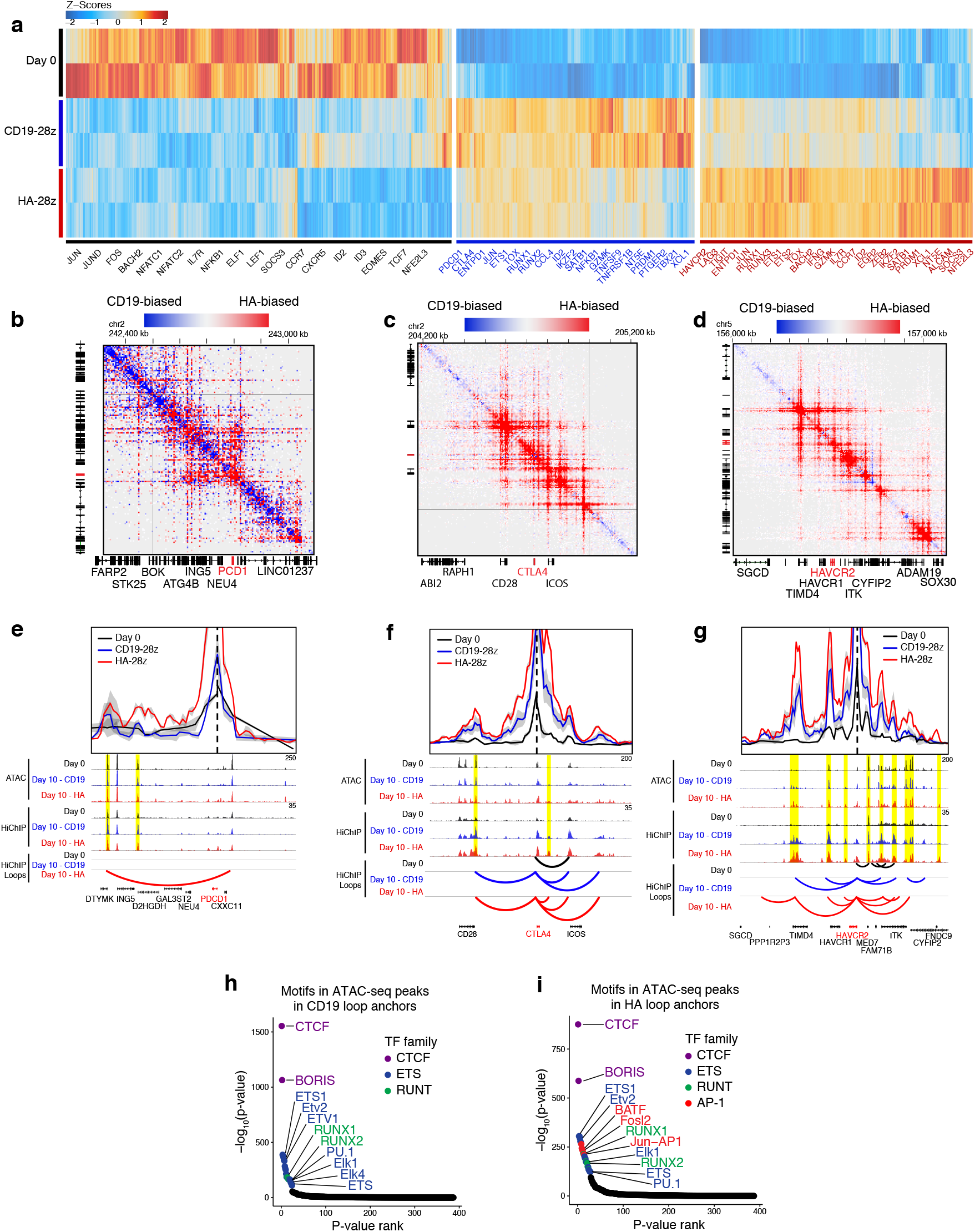
Gene-distal loci enriched in AP-1 TF motif sequences are differentially proximal to exhaustion-regulating genes. (A) The 5000 most variable 10-kb HiChIP loop anchors across Day 0 non-transfected primary T cells, Day 10 CD19-28z, and Day 10 HA-28z CAR T cells, clustered by interaction Z-scores and labeled with select genes whose transcription start sites lie within the loop anchor. (B) HiChIP interaction matrix of differentially proximal chromatin loci between Day 10 CD19-28z and HA-28z CAR T cells at the *PDCD1*, (C) *CTLA4*, and (D) *HAVCR2* loci. (E) Virtual-4C HiChIP interaction map anchored at the transcription start site and 1D sequencing coverage for both ATAC-seq and HiChIP at *PDCD1*, (F) *CTLA4*, and (G) *HAVCR2*. (H) The enrichment of TF motif binding sequences within each loop anchor, ranked by enrichment significance, in Day 10 CD19-28z and (I) HA-28z CAR T cells.

Interaction maps at genes associated with T cell exhaustion, such as *PDCD1, CTLA4*, and *HAVCR2*, demonstrate extensive distal chromosomal contact, often reaching hundreds of kilobases from the gene promoter (Figure 4b-d). The H3K27ac HiChIP signal is substantially stronger for these three loci associated with exhaustion in HA-28z than non-exhausted CD19 CAR T cells (Fig 4b-d). Virtual 4C analysis anchored at gene promoters highlights loci of differential chromosomal proximity that show both differential and similar levels of chromatin accessibility as measured by ATAC-seq (Figure 4e-g), indicating that 3D chromosome looping provides an additional level of control beyond enhancer accessibility. Transcription factor motif enrichment of accessible chromatin identified in ATAC-seq peaks are independently enriched in HiChIP loop anchors and identified the AP-1 family motif as specifically enriched in the exhausted HA-28z CAR T cell loop anchors (Figure 5h-i). In contrast, the motifs of the general DNA looping factors CTCF and its homolog BORIS, involved in enhancer-promoter contacts^47,48^, are equally enriched in HiChIP anchor points in HA-28z and CD19-28z CAR T cells (Figure 5h-i). These results suggest that AP-1 family factors likely drive both the chromatin accessibility and higher order 3D genome architecture for T cell exhaustion.

**Figure 5.**
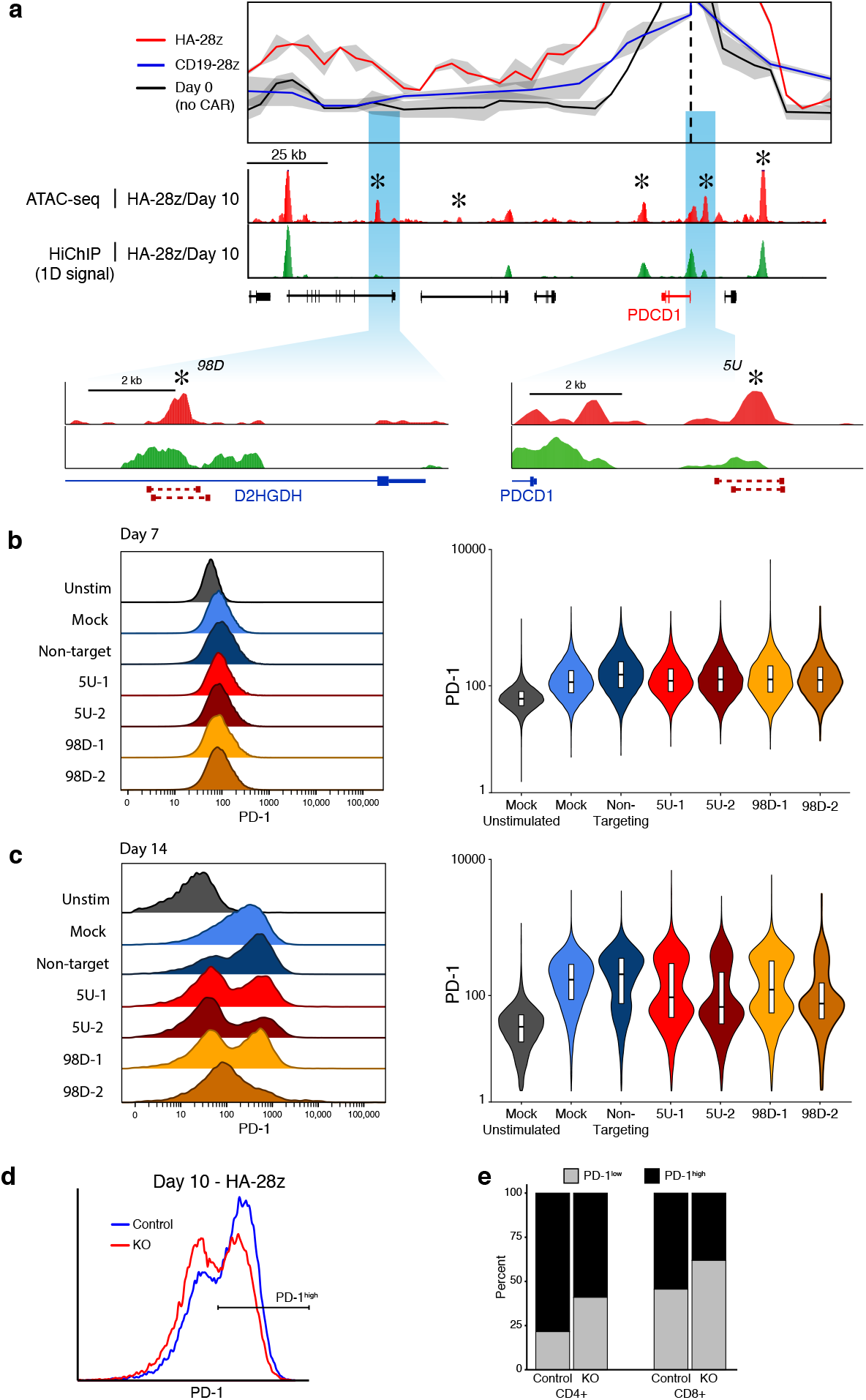
Genome editing at putative enhancer loci causes reduced expression of PD-1 in Jurkat cells. (A) Virtual-4C interaction map and 1D ATAC-seq and HiChIP sequencing coverage at the PDCD1 locus in Day 10 CD19-28z and HA-28z CAR T cells highlighting proximal (5 kb upstream) and distal (98 kb downstream) targets for CRISPR/Cas9 excision. (B) PD-1 expression seven days following stimulation with CD3/CD28 beads in unstimulated Jurkat cells, mock-transfected cells, Jurkat cells transfected with non-targeting Cas9-RNPs, and Jurkat cells transfected with Cas9-RNPs targeting the 5 kb-upstream or 98 kb-downstream regulatory loci. (C) PD-1 expression following 14 days of stimulation. (D) PD-1 expression in CD19-28z and HA-28z CAR T cells following 10 days of *ex vivo* culture and transfection with Cas9-RNPs targeting the 5 kb-upstream regulatory locus for excision. (E) The proportion of PD-1^high^ and PD-1^low^ HA-28z CAR T cells following Cas9-RNP transfection and non-targeting Cas9 transfection, broken down by CD4+/CD8+ T cell type.

### Genome editing of regulatory loci reduces expression of PD-1

To support our identification of regulatory loci using multi-omic assays, we used CRISPR-Cas9 gene editing in immortalized Jurkat T cells to remove putative regulatory sequences from the genome. We selected two loci near *PDCD1*, one 5 kb upstream and one 98 kb downstream of the transcriptional start site, each exhibiting differential chromatin accessibility between CD19-28z and HA-28z CAR T cells during the exhaustion time course, as well as 3D proximity with the promoter as reported by HiChIP (Figure 4e, 5a). Both loci have been previously demonstrated to be differentially accessible in the Jurkat T cell line following prolonged *in vitro* stimulation, a similarly dysfunctional phenotypic state to canonical *in vivo* T cell exhaustion (Supplemental Figure 5a-b)^49–51^. CRISPR-Cas9 guide RNA pairs were designed to flank the ATAC-seq peak at each locus, removing 700-1000 bp (Figure 5a).

We transfected Jurkat cells with purified guide RNA-Cas9 ribonucleoprotein (RNP) pairs targeting one of the exhaustion-associated peaks, purified RNP incorporating a non-targeting control guide RNA, or a mock-transfection without RNP. Following seven days of stimulation, all samples were consistently PD-1^low^, at levels similar to unstimulated Jurkat cells (Figure 5b, Supplemental Figure 5c). Following 14 days of stimulation, the mock and non-targeting RNP transfections resulted in cells predominantly PD-1^high^. In contrast, PD-1 expression was bimodal in cells that received RNPs targeting the exhaustion-associated enhancers, and the majority of cells remained PD-1^low^ (Figure 5c, Supplemental Figure 5d). Bimodality of PD-1 expression is likely due to inefficiency of CRISPR/Cas9 transfection and editing combined with incomplete penetrance of ablating single enhancers.

We then targeted the exhaustion-associated locus 5 kb-upstream of *PDCD1* for deletion in HA-28z CAR T cells. The CAR T cells were transfected with pairs of guide RNA-Cas9 RNPs concurrent with CAR transfection. Following 10 days of *ex vivo* culture, a greater proportion of cells were PD-1^low^, compared to CAR-only HA-28z CAR T cells (Figure 5d). Both CD4+ and CD8+ CAR T cells demonstrated an increased fraction of PD-1^low^ when the 5 kb-upstream enhancer was targeted for editing (Figure 5e). Together, these results demonstrate the feasibility of targeting exhaustion specific enhancers in human CAR T cells to potentially manipulate the exhaustion program.

## DISCUSSION

Here we describe the integrated landscape of accessible chromatin, histone acetylation, and long-range chromatin looping in a dynamic model of human CAR T cell exhaustion. These data extend our previously reported transcriptional and epigenetic features that differentiate non-exhausted CAR T cells and CAR T cells that have acquired the functional hallmarks of exhaustion^7^. Assaying the developing CAR T cells through a time course of their maturation, we uncovered the epigenetic and transcriptional regulatory features that mark different stages in the acquisition of the exhausted phenotype in HA-28z CAR T cells, as compared to non-exhausting CD19-28z CAR T cells. Exhausted CAR T cells show global alterations in their chromatin accessibility profiles at Day 7, which precedes the manifestation of phenotypic and functional hallmarks of exhaustion observed later in the time course. The global patterns of exhaustion-associated ATAC-seq peaks and the underlying TF motifs we identified are very similar across CD4+/CD8+ and naïve/central memory T cell subtypes (Figure 2b,d), suggesting the mechanisms we describe are general to broader T cell types.

We found putative enhancer elements and transcription factors such as NFE2L1 associated with the exhausted T cells at the time point preceding significant phenotypic divergence between CD19-28z and HA-28z CAR T cells, indicating their functional role in the development of exhaustion, perhaps setting up a regulatory landscape more conducive to a subsequent transcriptional program associated with the exhausted phenotype. NFE2L1, a bZIP family TF, has been previously shown to exhibit increased expression in activated primary T lymphocytes^52^ and association with the expression, protein abundance, and nuclear binding activity of several AP-1 factors^53^, including c-Jun, a TF we previously identified as a mediator of dysfunction in exhausted CAR T cells^7^. Our previously reported single-cell RNA-seq analysis of exhausted GD2-28z CAR T cells^7^ demonstrated that NFE2L1 and its homolog NFE2L3 are more highly expressed in a GD2-28z-specific subset of cells that also express exhaustion markers PD-1, CTLA4, and LAG3 (Supplementary Figure 6a-b).

When comparing the exhausted human CAR T cell chromatin state to published mouse models of T cell exhaustion, we identified distinct chromatin signatures marking exhaustion in each species. The majority of accessible chromatin loci we found in exhausted human CAR T cells were not accessible in dysfunctional mouse T cells, either because the locus in mouse was not accessible or there was no homologous sequence in mouse (and vice versa). Even though much of the dysfunctional chromatin signature was unique to each species, the transcription factor motif enrichment and associated-gene pathway enrichment for those signatures were largely shared.

Cancer immunotherapies must find a balance between mounting a strong antitumor immune response and overstimulation of the immune system. Immune checkpoint inhibitors, namely anti-PD-1 and anti-PD-L1 therapeutics, have been associated with cytokine-release syndrome^54^, a potentially fatal condition characterized by hyperactivation and massive release of cytokines by the immune system^55^. CD19-targeting CAR T cells have themselves been associated with cytokine-release syndrome in patients^2,56^. One challenge lies in tuning the expression levels of key regulators of dysfunction during specific cell states to prevent dysfunction while maintaining robust anti-tumor activity. A notable example is the *Tox* gene, found to be a driver of exhaustion in a mouse model of T cell exhaustion, yet deletion of the gene results in poor T cell persistence in tumors^57^.

Enhancers are a promising target for genome editing in the search for effective treatment for genetic diseases^58,59^. The benefits of editing enhancer elements, rather than gene deletion or other methods of editing the gene promoter or body, are twofold: enhancers are cell type-specific and gene expression remains possible under different cell states under the control of different enhancer elements. We found that deletion of *PDCD*1-assocaited enhancers reduced the expression of PD-1 in an *in vitro* model of T cell exhaustion. The enhancers were identified using a combination of chromatin accessibility profiling, chromosome conformation capture, and transcriptome analysis, highlighting the importance of multi-omic profiling in order to capture a more full picture of the genetic regulatory networks. For example, one exhaustion-biased enhancer of *PDCD1* that we identified was 98 kb from the *PDCD*1 promoter, spanning a region that includes several genes and residing within another gene’s intron. Many current methods of enhancer analysis rely on nearest-gene associations to assign regulatory function, but evidence is mounting that this is insufficient without including orthogonal techniques, such as chromosome conformation.

We were also able to delete the exhaustion-associated enhancer most proximal to the *PDCD1* promoter in HA-28z CAR T cells, and we found a marked reduction in PD-1^high^ CAR T cells in the population following *ex vivo* maturation. This suggests that the epigenetic landscape of CAR T cells may be modulated to improve their efficacy prior to reinfusion into patients. Importantly, the *PDCD1* gene itself was not deleted, leaving open the possibility of its expression in non-exhaustion contexts and cell states. More investigation is required to explore the functional consequences and potential of the deleted enhancer, especially in regard to the cells’ antitumor response, their *in vivo* ability to avoid exhaustion, and the prevention of overstimulation and cytokine-storm syndrome.

## MATERIALS AND METHODS

### T cell isolation

Buffy coats from healthy donors were obtained from the Stanford Blood Center. T cells were purified using the RosetteSep Human T Cell Enrichment Kit (Stem Cell Technologies) with the Lymphoprep density gradient medium and SepMate-50 tubes. 2×10^7^ T cells were aliquoted into CryoStor cryopreservation media (Stem Cell technologies) for long-term storage.

### CAR Vector Production

The 293GP cell line was used to produce MSGV retroviral vectors encoding CD19-28z and HA-28z CARs, as previously described. At 24 and 48 hours following plasmid transfection, the culture media was replaced. Supernatants were collected and replaced at 48 and 72 hours following transfection. Cell debris was removed by centrifugation, and the supernatants frozen at −80°C.

### CAR T Cell Production

Primary T cells were thawed and cultured in AIM V media + 5% FBS, 10mM HEPES, 2mM GlutaMAX, 100 U/mL penicillin, and 10 *μ*g/mL streptomycin (Gibco). T cells were activated the same day using Human T-Expander CD3/CD28 Dynabeads (Gibco) in a 3:1 beads:cells ratio and 100 U/mL recombinant human IIL-2 (Peprotech). Non-tissue culture treated 12-well plates were prepared by incubating 1 mL PBS + 25 *μ*g/mL Retronectin (Takara) overnight at 4°C. Wells were subsequently washed with PBS and blocked with 2% BSA for 15 minutes. 1 mL thawed retroviral supernatant was added to each well and centrifuged for 2 hours at 3,200 RPM at 32°C. On day 2 following T cell activation, cells were added to the prepared plates and maintained at 0.5-1×10^6^ cell/mL.

### Flow Cytometry Antibodies

BioLegend: CD4-APC-Cy7 (clone OKT4), CD8-PerCp-Cy5.5 (clone SK1), TIM-3-BV510 (clone F38-2E2), CD39-FITC or APC-Cy7 (clone A1), CD95-PE (clone DX2), CD3-PacBlue (clone HIT3a), PD-1-BV421 (clone EH12.2H7), CD3-PE (clone HIT3a)
eBioscience: PD-1-PE-Cy7 (clone eBio J105), LAG-3-PE (clone 3DS223H), CD45RO-PE-Cy7 (clone UCHL1), CD45-PerCp-Cy5.5 (clone HI30)
BD: CD45RA-FITC or BV711 (clone HI100), CCR7-BV421 (clone 150503), CD122-BV510 (clone Mik-β3), CD62L-BV605 (clone DREG-56), CD4-BUV395 (clone SK3), CD8-BUV805 (clone SK1)
Anti-CD19-28z CAR antibody was provided by B. Jena and L. Cooper. The anti-HA-28z CAR antibody was provided by NCI-Frederick. Each was conjugated with Dylight288 and/or 650 antibody labelling kits (Thermo Fisher).

### Cytokine Production

1×10^5^ CAR^+^ T cells were cultured for 24 hours in 96-well flat-bottom plates in 200 *μ*L media in triplicate. ELISA (BioLegend) for IFNγ and IL-2 was conducted on collected supernatants.

### Omni-ATAC-seq

Omni-ATAC-seq was conducted as previously described. Briefly, 100,000 CAR T cells were sorted into complete media, centrifuged at 4°C at 500 g, and resuspended in ATAC-seq resuspension buffer (RSB) (10 mM Tris-HCl, 10 mM NaCl, 3 mM MgCl2, 0.1% NP-40, 0.1% Tween-20, and 0.01% digitonin). Cell suspensions were divided into two replicates. Samples were placed on ice for 3 minutes and washed with RSB-2 (10 mM Tris-HCl, 10 mM NaCl, 3 mM MgCl2, and 0.1% Tween-20). Centrifugation at 500 g for 10 minutes at 4°C made a pellet of nuclei, which was resuspended in 50 *μ*L transposition mix (25 μL 2× TD buffer, 2.5 μL transposase (Illumina), 16.5 μL PBS, 0.5 μL 1% digitonin, 0.5 μL 10% Tween-20, 5 μL H2O). Transposition was conducted at 37°C for 30 minutes in a shaking thermomixer at 1000 RPM. DNA was purified with the MinElute PCR Purification Kit (Qiagen). Libraries were amplified by PCR using the NEBNext Hi-Fidelity PCR Master Mix and custom primers (IDT) and quantified with the KAPA Library Quantification Kit. Libraries were sequenced on the Illumina NextSeq at the Stanford Functional Genomics Facility with paired-end 75-bp reads.

Adapter sequences were trimmed from the reads and sequences aligned to the hg19 genome with bowtie2. Reads were filtered with Picard tools against mitochondrial sequences, quality scores <20, and PCR duplicates. Peaks were called using MACS2 on Tn5-corrected insertion sites. A union peak set was compiled by extending peak summits to 500 bp, merging all summits, running bedtools cluster, selecting summits with the highest MACS2 score, and filtering b the ENCODE hg19 blacklist. Peaks were annotated with HOMER^60^. TF motifs were found by chromVAR^61^ motifmatchr with the chromVARmotifs HOMER set and HOMER. Genome coverage tracks were made by pileups binning every 100 bp with rtracklayer. Differential peaks calling was done with DESeq2^62^ using TSS-annotated peaks as control loci or consistent ‘housekeeping peaks.’ Clustering was performed using the regularized-log transform values from DESeq2. TF motif enrichment was found using a hypergeometric test on the motifmatchr representation in peak subsets.

### RNA-seq

Total RNA was collected from 2×10^6^ CAR T cells with the RNEasy Plus Mini isolation kit (Qiagen). Library preparation and RNA-seq was performed by BGI America (Cambridge, MA) on the BGISEQ-500 as single-end 50-bp reads.

### Human and mouse ATAC-seq data comparison

Mouse ATAC-seq fastq files were downloaded from GEO (GSM2337202)^21^. Mouse ATAC-seq data were aligned and filtered based on the mm9 reference genome, and were analyzed with the same pipeline and tools as human Omni-ATAC-seq. Peaks within ±3000bp were categorized as promoter, peaks that overlap with the Fantom5^63^ CD4+ T cell enhancer list were defined as enhancer. Each ATAC-seq peak was annotated with its nearby genes using GREAT^64^ by basal plus extension default setting. The UCSC tools liftOver module was used to lift peaks over between mm9 and hg19. Bedtools intersect module with default settings were used to define the species-specific peaks and conservative peaks. Peaks that cannot be lifted over between mm9 and hg19 were defined as species-specific peaks without homology. Mouse peaks which could be lifted over from mm9 to hg19, but do not overlap with any human peaks, were defined as mouse-specific peaks with homology. Human-specific peaks with homology were defined with the same method. Conserved peaks were peaks that could be lifted over, and the corresponding genomic region is accessible in both human and mouse samples. The MultiBamCov module from bedtools was used to generate peak read count matrix from bam files. Accessibility Z-scores of motifs were generated by the chromVar R package using Jaspar motifs, and the top motifs with the highest variability were reported. DESeq2 was used to call differentially accessible peaks between exhausted T cells and non-exhausted T cells at day 7 in human samples, and day 8 in mouse samples, with FDR < 0.05 and fold-change of >2. Motif enrichment analysis based on the differential ATAC-seq peaks was conducted by Homer with FDR < 0.05.

### HiChIP

500,000 cells per replicate were pelleted at 300 *g* for 5 minutes and resuspended in 500 μL 1% methanol-free formaldehyde (Thermo Fisher). After 10 minutes room temperature incubation with rotation, cross-linking was quenched with glycine to a final concentration of 125 mM. Cross-linked cells were washed in PBS and pellets stored at −80 °C until subsequent next steps. Pellets were resuspended in 500 *μ*L cold Hi-C lysis buffer (10 mM Tris-HCl pH 7.5, 10 mM NaCl, 0.2% NP-40, 1X protease inhibitor (Roche)) and rotated for 30 minutes at 4 °C. Samples were centrifuged at 2500 *g* for 5 minutes at 4 °C to pellet nuclei. Pelleted nuclei were washed in cold Hi-C lysis buffer and resuspended in 100 μL of 0.5% SDS at 62 °C for 10 minutes without rotation. SDS was quenched by adding 285 μL of water and 50 μL of 10% Triton X-100, with rotation at 37 °C for 30 minutes. Fragmentation was carried out by adding 50 *μ*L NEB Buffer 2 and 8 μL of 25 U/μL MboI restriction enzyme (New England Biolabs). Samples were incubated at 37 °C for 2 hours with rotation, then 62 °C for 20 minutes. 52 *μ*L of fill-in master mix were added, consisting of 37.5 0.4 mM biotin-dATP (Thermo Fisher), 1.5 *μ*L each 10 mM dCTP/dGTP/dTTP (NEB), and 10 *μ*L 5U/*μ*L DNA Pol I Large Klenow fragment (NEB), then incubated at 37 °C for 1 hour. 948 *μ*L ligation master mix were added, consisting of 150 *μ*L 10X NEB T4 DNA ligase buffer with 10 mM ATP (NEB), 125 *μ*L 10% Triton X-100, 3 *μ*L 50 mg/mL BSA, 10 *μ*L 400 U/*μ*L T4 DNA Ligase (NEB), and 660 *μ*L water, then incubated at room temperature for 4 hours. Nuclei were then pelleted at 2500 *g* for 5 minutes, supernatant discarded, and resuspended in 880 *μ*L nuclear lysis buffer (50 mM tris-HCl, 10 mM EDTA, 1% SDS, 1X protease inhibitor). DNA was sonicated in a Covaris E220 and Covaris millitube with the following parameters: fill level 10, duty cycle 5, PIP 140, cycles/burst 200, time 4 minutes. Samples were centrifuged for 15 minutes at 4 °C at 16,000 *g*. 2X volume of ChIP Dilution Buffer (0.01% SDS, 1.1% Triton X-100, 1.2 mM EDTA, 16.7 mM Tris-HCl pH 7.5, 167 mM NaCl) was added, and each total sample was split across two 1.5 mL tubes. 21 *μ*L Protein A beads (Thermo Fisher), for our total of 3.5 million cells, were washed in ChIP dilution buffer and resuspended in 50 *μ*L ChIP dilution buffer per tube (100 *μ*L total). Beads were added to sample tubes and incubated at 4 °C for 1 hour with rotation. Samples were positioned on a magnet, and the supernatant transferred to new tubes. 2.625 *μ*g anti-H3K27ac antibody (Abcam) were diluted 10X in ChIP dilution buffer (for 3.5 million cells total), distributed equally across tubes, and incubated overnight at 4 °C with rotation. Samples were put on a magnet and supernatants transferred to new tubes. 21 μL Protein A beads were washed in ChIP dilution buffer, resuspended in 50 μL ChIP dilution buffer per tube (100 μL total), added to each sample, and incubated for 2 hours at 4 °C with rotation. Samples were placed a magnet and the beads washed three times each with low-salt wash buffer (0.1% SDS, 1% Triton X-100, 2 mM EDTA, 20 mM Tris-HCl pH 7.5, 150 mM NaCl), high-salt wash buffer (0.1% SDS, 1% Triton X-100, 2 mM EDTA, 20 mM Tris-HCl pH 7.5, 500 mM NaCl), and LiCl wash buffer (10 mM Tris-HCl pH 7.5, 250 mM LiCl, 1% NP-40, 1% sodium deoxycholate, 1 mM EDTA). Beads were resuspended in 100 μL DNA Elution buffer (50 mM sodium bicarbonate pH 8.0, 1% SDS) and incubated at room temperature for 10 minutes with rotation, then at 37° C for 3 minutes with shaking. Samples were placed on a magnet and supernatants transferred to new tubes. Elution was repeated with another 100 *μ*L elution buffer, transferring to the same tube. 10 *μ*L Proteinase K (Thermo Fisher) were added to each sample and incubated at 55 °C for 45 minutes with Shaking, followed by 65 °C for 1.5 hours with shaking. Samples were purified with Zymo Research DNA Clean and Concentrator columns and eluted in 10 μL of water. 5 μL Streptavidin C-1 beads (Thermo Fisher) were Washed with Tween Wash Buffer (5 mM Tris-HCl pH 7.5, 0.5 mM EDTA, 1 M NaCl, 0.05% Tween-20) and resuspended in 10 μL 2X Biotin Binding Buffer (10 mM Tris-HCl pH 7.5, 1 mM EDTA, 2M NaCl). Beads were added to samples and incubated at room temperature for 15 minutes with rotation. Samples were placed on a magnet and supernatants discarded. Beads were washed twice in 500 *μ*L Tween wash buffer with 2 minutes shaking at 55 °C. Beads were washed in 100 μL 1X TD Buffer (2X TD Buffer: 20 mM Tris-HCl pH 7.5, 10 mM magnesium chloride, 20% dimethylformamide). Beads were resuspended in 25 μL 2X TD Buffer. 2.5 μL Tn5 transposase (Illumina) was added for each 50 ng of DNA in each sample, and water was added up to 50 *μ*L. Samples were incubated at 55° C for 10 minutes with shaking. Samples were placed on a magnet and supernatants removed. Beads were resuspended in 50 mM EDTA and incubated for 30 minutes at 50 °C. Samples were placed on a magnet and supernatants removed. Samples were washed twice more with 50 mM EDTA at 50 °C for 3 minutes. Samples were washed twice with Tween wash buffer at 55 °C for 2 minutes. Samples were washed in 10 mM Tris. Beads were resuspended in 50 *μ*L PCR master mix (25 *μ*L 2X Phusion HF buffer (NEB), 1 *μ*L 12.5 *μ*M Nextera Ad1 primer (Illumina), 1 *μ*L 12.5 *μ*M Nextera Ad2 barcoded primer (Illumina), 23 *μ*L water). PCR was run as follows: 72 °C for 5 minutes, 98 °C for 1 minute, and 8 cycles of 98 °C for 15 seconds, 63 °C for 30 seconds, and 72 °C for 1 minute. Amplified libraries were placed on a magnet and transferred to new tubes. Libraries were Zymo column-purified and eluted together into the same 10 *μ*L water. Libraries were PAGE purified and size-selected for 300-700 bp, then column-cleaned into 10 *μ*L water. Libraries were sequenced on an Illumina NextSeq at the Stanford Functional Genomics Facility with paired-end 75-bp reads.

HiChIP data was analyzed as previously described^47^. Paired-end reads were aligned to hg19 with the Hi-C Pro pipeline, interactions called with Juicer HICCUPS^65^ and Fit-HiC^66^, and interaction maps made with Juicebox^67^.

### Cell Lines

The Jurkat T cell line was obtained from ATCC (clone E6-1) and cultured in RPMI + 10% FBS, 10 mM HEPES, 100 U/mL penicillin, and 100 *μ*g/mL streptomycin (Gibco). Cells were stimulated with Human T-Expander CD3/CD28 Dynabeads (Gibco). At sorting, propidium iodide was used to stain and exclude dead and dying cells.

### CRISPR/Cas9 editing

Cas9-ribonucleoprotein (RNP) complexes were delivered into Jurkat cells using the Alt-R CRISPR-Cas9 system (IDT) and the Neon Transfection System (Thermo Fisher). Guide-RNA sequences were designed using the Broad Institute’s online sgRNA design tool and CRISPETa paired guide designer^68^. RNA oligonucleotides were synthesized by IDT:

PDCD1 -5kb-upstream-1-5’: /AltR1/rGrUrArUrUrGrCrCrArArGrGrGrCrArCrCrCrGrGrGrUrUrUrUrArGrArGrCrUrArUrGr CrU/AltR2/
PDCD1 -5kb-upstream-1-3’: /AltR1/rUrCrGrUrUrGrGrGrCrUrGrUrUrUrGrUrUrArArGrGrUrUrUrUrArGrArGrCrUrArUrG rCrU/AltR2/
PDCD 1 -5kb-upstream-2-5’: /AltR1/rArUrGrCrGrUrArCrUrGrCrArCrCrUrUrCrCrCrArGrUrUrUrUrArGrArGrCrUrArUrGr CrU/AltR2/
PDCD 1 -5kb-upstream-2-3’: /AltR1/rArArUrCrGrUrUrGrGrGrCrUrGrUrUrUrGrUrUrArGrUrUrUrUrArGrArGrCrUrArUrG rCrU/AltR2/
PDCD 1 -98kb-downstream-1-5’: /AltR1/rArUrCrUrGrUrCrUrGrCrUrArArGrGrUrCrCrArCrGrUrUrUrUrArGrArGrCrUrArUrGr CrU/AltR2/
PDCD 1 -98kb-downstream-1-3’: /AltR1/rArUrGrArArCrArUrArUrUrCrUrGrUrGrGrArUrGrGrUrUrUrUrArGrArGrCrUrArUrG rCrU/AltR2/
PDCD 1 -98kb-downstream-2-5’: /AltR1/rArUrCrUrGrArCrGrGrCrCrGrArUrCrArCrArCrArGrUrUrUrUrArGrArGrCrUrArUrGr CrU/AltR2/
PDCD 1 -98kb-downstream-2-3’: /AltR1/rArCrGrArUrCrArCrUrCrUrArCrUrGrCrCrCrUrCrGrUrUrUrUrArGrArGrCrUrArUrGr CrU/AltR2/

Pairs of RNPs were assembled and transfected into Jurkat cells according to manufacturer’s protocol. Briefly, 220 pmol of each guide-RNA in a pair and 440 pmol tracrRNA were combined in 10 *μ*L TE buffer, heated to 95° for 5 min, then placed at room temperature to cool. 10 *μ*L of 36 *μ*M Cas9 enzyme was added to each crRNA-tracrRNA pair and incubated at room temperature for 10 minutes. For each transfection, 5×10^6^ Jurkat cells were collected, centrifuged, washed in PBS, and resuspended in 80 *μ*L Buffer R. The 80 *μ*L cell suspension was mixed with the 20 *μ*L RNP complex and electroporated in the Neon Transfection System at 1600 V, 10 ms pulse width, and 3 pulses. Cells were transferred to pre-warmed media in 6-well plates. At 24 hours, cells were stimulated with Human T-Expander CD3/CD28 Dynabeads (Gibco) in a 3:1 beads:cells ratio and 100 U/mL recombinant human IL-2 (Peprotech).

## ACKNOWLEDGEMENT

We thank members of our labs for discussion. Supported by Parker Institute for Cancer Immunotherapy (A.T.S., C.L.M., H.Y.C.), NIH RM1-HG007735 (W.J.G., W.W., H.Y.C.), and NIH K08CA230188 (A.T.S.). H.Y.C. is an Investigator of the Howard Hughes Medical Institute. E.W.W. was supported by a Cellular and Molecular Immunobiology Training Grant (5 T32 AI07290, NIH NIAID, E.W.W.)

