## Supplemental Figures for "Dynamic chromatin regulatory landscape of human CAR T cell exhaustion"

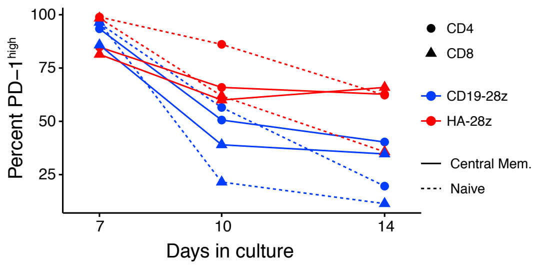


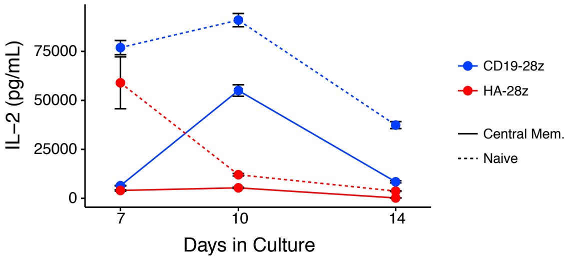


**c**

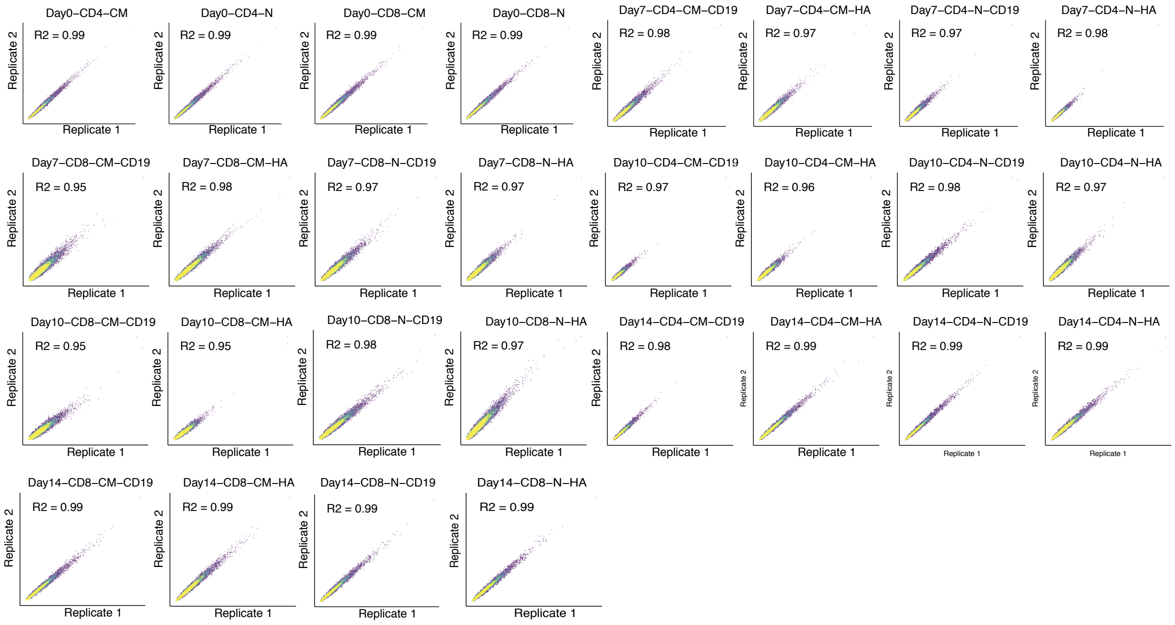


**d**

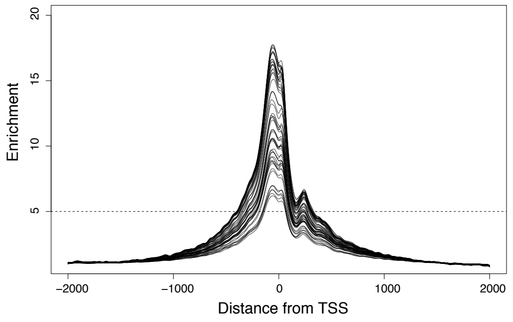


**Supplemental Figure 1. Donor-derived CAR T cell phenotyping and Omni-ATAC-seq quality control metrics.** (A-B) Surface expression and cytokine secretion of markers of T cell exhaustion throughout the CAR T cell maturation time course. (C) All samples demonstrate high replicate concordance. (D) All samples demonstrate high enrichment of genome alignment to transcription start sites.

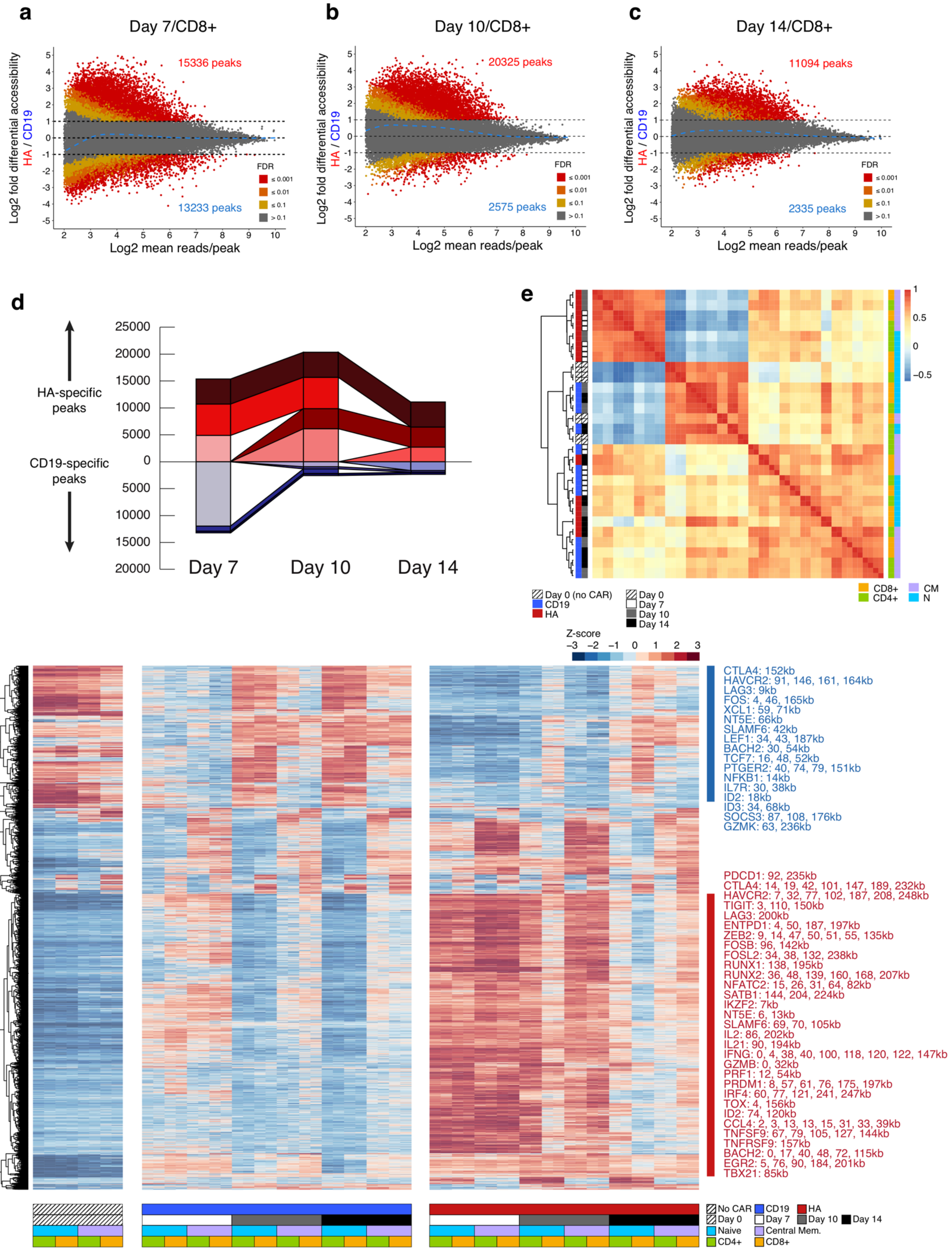
a
Day 7/CD8+
15336 peaks
13233 peaks
b
Day 10/CD8+
20325 peaks
2575 peaks
c
Day 14/CD8+
11094 peaks
2335 peaks
Log2 fold differential accessibility
HA / CD19
Log2 mean reads/peak
FDR
≤ 0.001
≤ 0.01
≤ 0.1
> 0.1
d
HA-specific peaks
CD19-specific peaks
Day 7
Day 10
Day 14
e
Day 0 (no CAR)
CD19
HA
Day 0
Day 7
Day 10
Day 14
CD8+
CD4+
CM
N
f
Z-score
CTLA4: 152kb
HAVCR2: 91, 146, 161, 164kb
LAG3: 9kb
FOS: 4, 46, 165kb
XCL1: 59, 71kb
NT5E: 66kb
SLAMF6: 42kb
LEF1: 34, 43, 187kb
BACH2: 30, 54kb
TCF7: 16, 48, 52kb
PTGER2: 40, 74, 79, 151kb
NFKB1: 14kb
IL7R: 30, 38kb
ID2: 18kb
ID3: 34, 68kb
SOCS3: 87, 108, 176kb
GZMK: 63, 236kb
PDCD1: 92, 235kb
CTLA4: 14, 19, 42, 101, 147, 189, 232kb
HAVCR2: 7, 32, 77, 102, 187, 208, 248kb
TIGIT: 3, 110, 150kb
LAG3: 200kb
ENTPD1: 4, 50, 187, 197kb
ZEB2: 9, 14, 47, 50, 51, 55, 135kb
FOSB: 96, 142kb
FOSL2: 34, 38, 132, 238kb
RUNX1: 138, 195kb
RUNX2: 36, 48, 139, 160, 168, 207kb
NFATC2: 15, 26, 31, 64, 82kb
SATB1: 144, 204, 224kb
IKZF2: 7kb
NT5E: 6, 13kb
SLAMF6: 69, 70, 105kb
IL2: 86, 202kb
IL21: 90, 194kb
IFNG: 0, 4, 38, 40, 100, 118, 120, 122, 147kb
GZMB: 0, 32kb
PRF1: 12, 54kb
PRDM1: 8, 57, 61, 76, 175, 197kb
IRF4: 60, 77, 121, 241, 247kb
TOX: 4, 156kb
ID2: 74, 120kb
CCL4: 2, 3, 13, 13, 15, 31, 33, 39kb
TNFSF9: 67, 79, 105, 127, 144kb
TNFRSF9: 157kb
BACH2: 0, 17, 40, 48, 72, 115kb
EGR2: 5, 76, 90, 184, 201kb
TBX21: 85kb
No CAR
CD19
HA
Day 0
Day 7
Day 10
Day 14
Naive
Central Mem.
CD4+
CD8+

**Supplemental Figure 2. The dynamics of accessible chromatin loci over time identify loci with varying temporal control.** (A-C) Differentially accessible chromatin loci (peaks) in CD8+ CAR T cells at 7, 10, and 14 days following CAR transfection. (D) Differentially accessible loci across time points, colored by which timepoints and the number of timepoints in which they are differentially accessible. Loci differentially accessible in HA-28z CAR T cells (red) are more numerous persist through multiple time points to a greater degree and are more numerous than the loci differentially accessible in CD19-28z CAR T cells (blue). (E) Sample clustering by Pearson correlation of global chromatin accessibility profiles. (F) Top 5000 most variable chromatin accessibility peaks across all samples colored by z-score of peak accessibility in each sample. Many of the peaks differentially accessible in either the CD19-28z or HA-28z CAR T cells appear proximal to known exhaustion-associated and T cell effector genes. Distances between peaks and select gene promoters are listed.

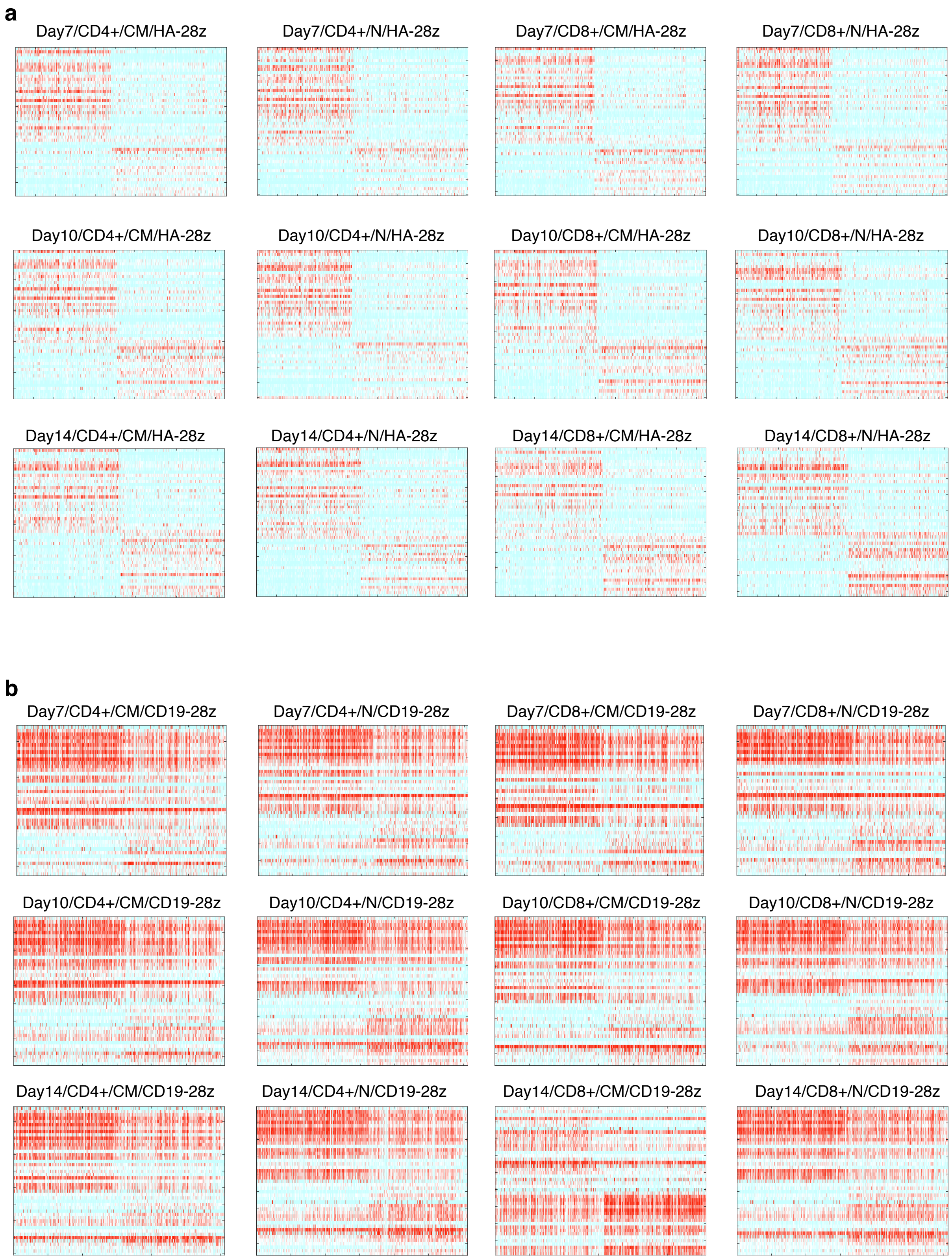


**Supplemental Figure 3. Transcription factor-gene dynamics demonstrate coregulated modules in exhausted CAR T cells.** PECA2 coregulation score heatmaps for all (A) HA-28z and (B) CD19-28z CAR T cell samples, merging replicates, indicating the strength of correlation between transcription and accessibility dynamics across samples.

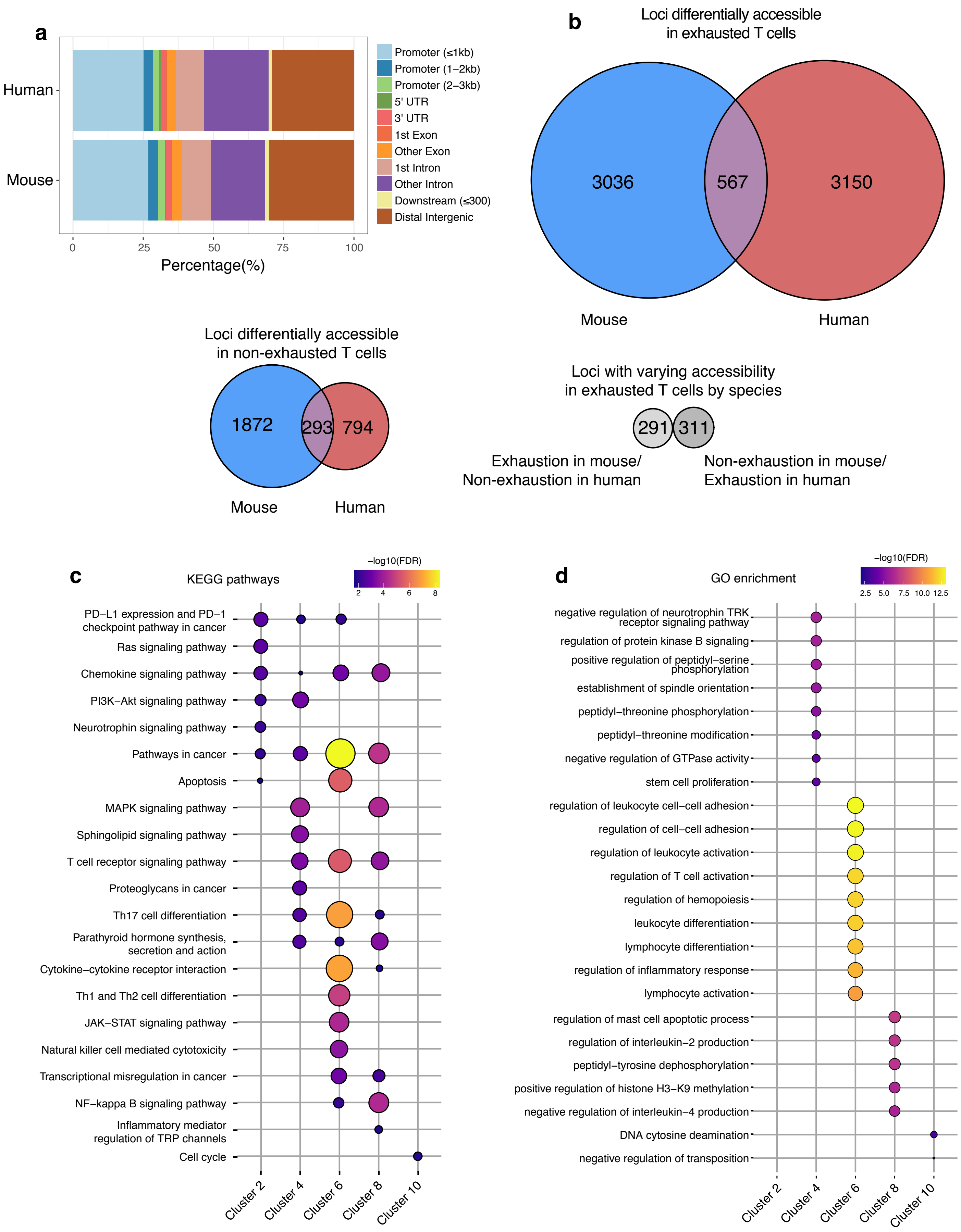

a
Human
Mouse
0
25
50
75
100
Percentage(%)
Promoter (≤1kb)
Promoter (1–2kb)
Promoter (2–3kb)
5' UTR
3' UTR
1st Exon
Other Exon
1st Intron
Other Intron
Downstream (≤300)
Distal Intergenic
b
Loci differentially accessible in exhausted T cells
3036
567
3150
Mouse
Human
Loci differentially accessible in non-exhausted T cells
1872
293
794
Mouse
Human
Loci with varying accessibility in exhausted T cells by species
291
311
Exhaustion in mouse/ Non-exhaustion in human
Non-exhaustion in mouse/ Exhaustion in human
c
KEGG pathways
−log10(FDR)
2
4
6
8
PD−L1 expression and PD−1 checkpoint pathway in cancer
Ras signaling pathway
Chemokine signaling pathway
PI3K−Akt signaling pathway
Neurotrophin signaling pathway
Pathways in cancer
Apoptosis
MAPK signaling pathway
Sphingolipid signaling pathway
T cell receptor signaling pathway
Proteoglycans in cancer
Th17 cell differentiation
Parathyroid hormone synthesis, secretion and action
Cytokine−cytokine receptor interaction
Th1 and Th2 cell differentiation
JAK−STAT signaling pathway
Natural killer cell mediated cytotoxicity
Transcriptional misregulation in cancer
NF−kappa B signaling pathway
Inflammatory mediator regulation of TRP channels
Cell cycle
Cluster 2
Cluster 4
Cluster 6
Cluster 8
Cluster 10
d
GO enrichment
−log10(FDR)
2.5
5.0
7.5
10.0
12.5
negative regulation of neurotrophin TRK receptor signaling pathway
regulation of protein kinase B signaling
positive regulation of peptidyl−serine phosphorylation
establishment of spindle orientation
peptidyl−threonine phosphorylation
peptidyl−threonine modification
negative regulation of GTPase activity
stem cell proliferation
regulation of leukocyte cell−cell adhesion
regulation of cell−cell adhesion
regulation of leukocyte activation
regulation of T cell activation
regulation of hemopoiesis
leukocyte differentiation
lymphocyte differentiation
regulation of inflammatory response
lymphocyte activation
regulation of mast cell apoptotic process
regulation of interleukin−2 production
peptidyl−tyrosine dephosphorylation
positive regulation of histone H3−K9 methylation
negative regulation of interleukin−4 production
DNA cytosine deamination
negative regulation of transposition
Cluster 2
Cluster 4
Cluster 6
Cluster 8
Cluster 10

**Supplemental Figure 4. Features of human and mouse dysfunctional T cell accessible chromatin.** (A) Genomic features of the accessible chromatin loci in exhausted human HA-28z CAR T cells and chronically stimulated mouse T cells (from Sen et al., 2016). (B) Quantification of the ATAC-seq peaks in the T cells of each species associated with (1) exhausted T cells, (2) non-exhausted T cells, or (3) both exhausted and non-exhausted T cells with varying accessibility by species. (C) KEGG pathway enrichment and (D) Gene Ontology (GO) term enrichment for genes associated with the ATAC-seq peaks within clusters of exhaustion-accessible ATAC-seq peaks in mouse and/or human T cells.

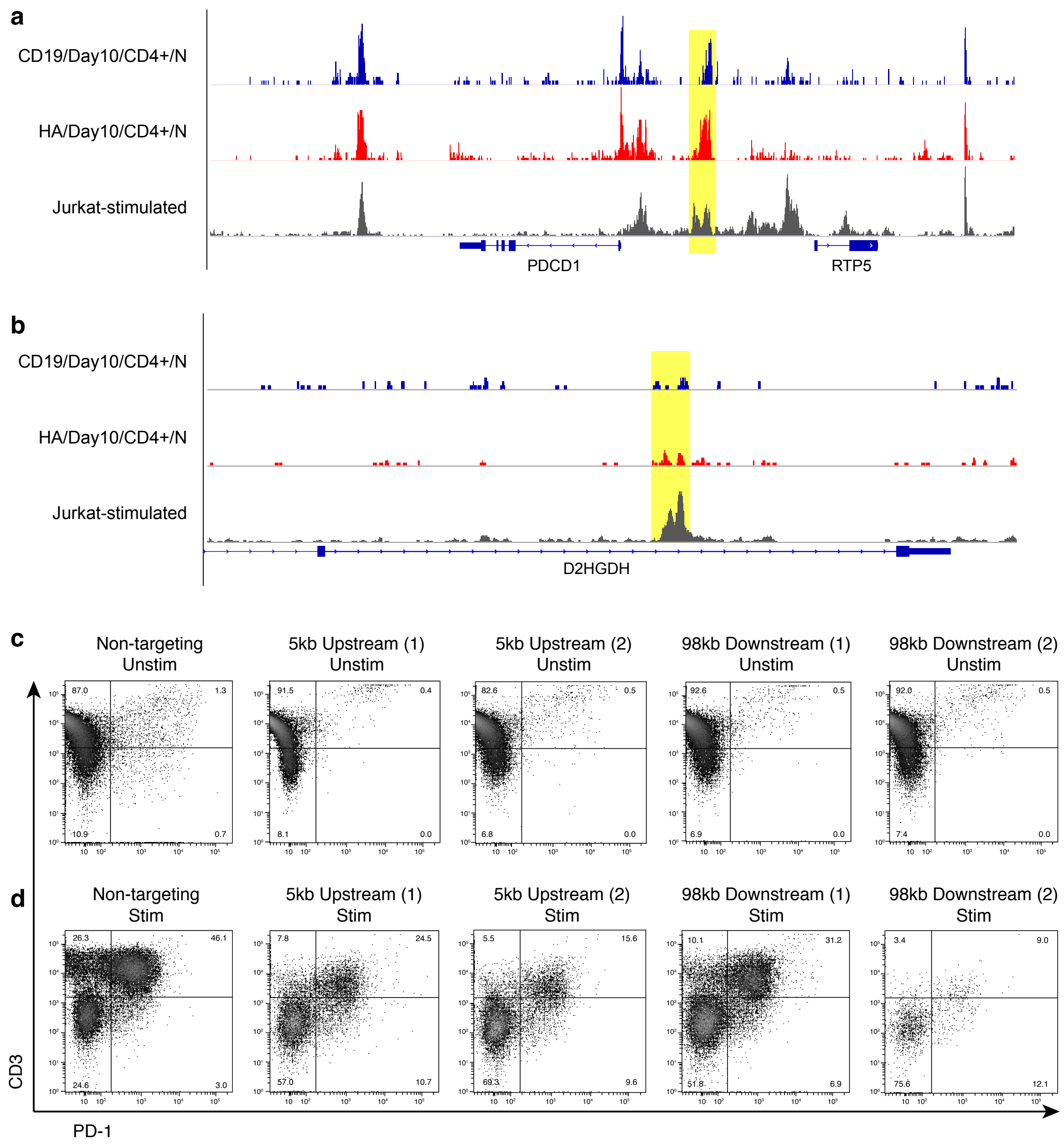


**Supplemental Figure 5. The Jurkat T cell line shares regulatory loci with stimulated and exhausted CAR T cells.** (A) Accessible chromatin sequencing alignment tracks of Day 10 CD19-28z, Day 10 HA-28z CAR T cells, and stimulated Jurkat cells (from Brignall et al., 2017) at the PDCD1 promoter locus and (B) 98 kb downstream from the PDCD1 transcription start site. (C) Cell surface expression of CD3 and PD-1 in unstimulated and (D) IL-2/CD3/CD28-stimulated Jurkat cells treated with non-targeting CRISPR/Cas9 RNPs, RNP pairs targeting the exhaustion-associated accessible chromatin locus 5 kb upstream of the PDCD1 promoter, or RNP pairs targeting 98 kb downstream of PDCD1 following 14 days of *in vitro* culture.

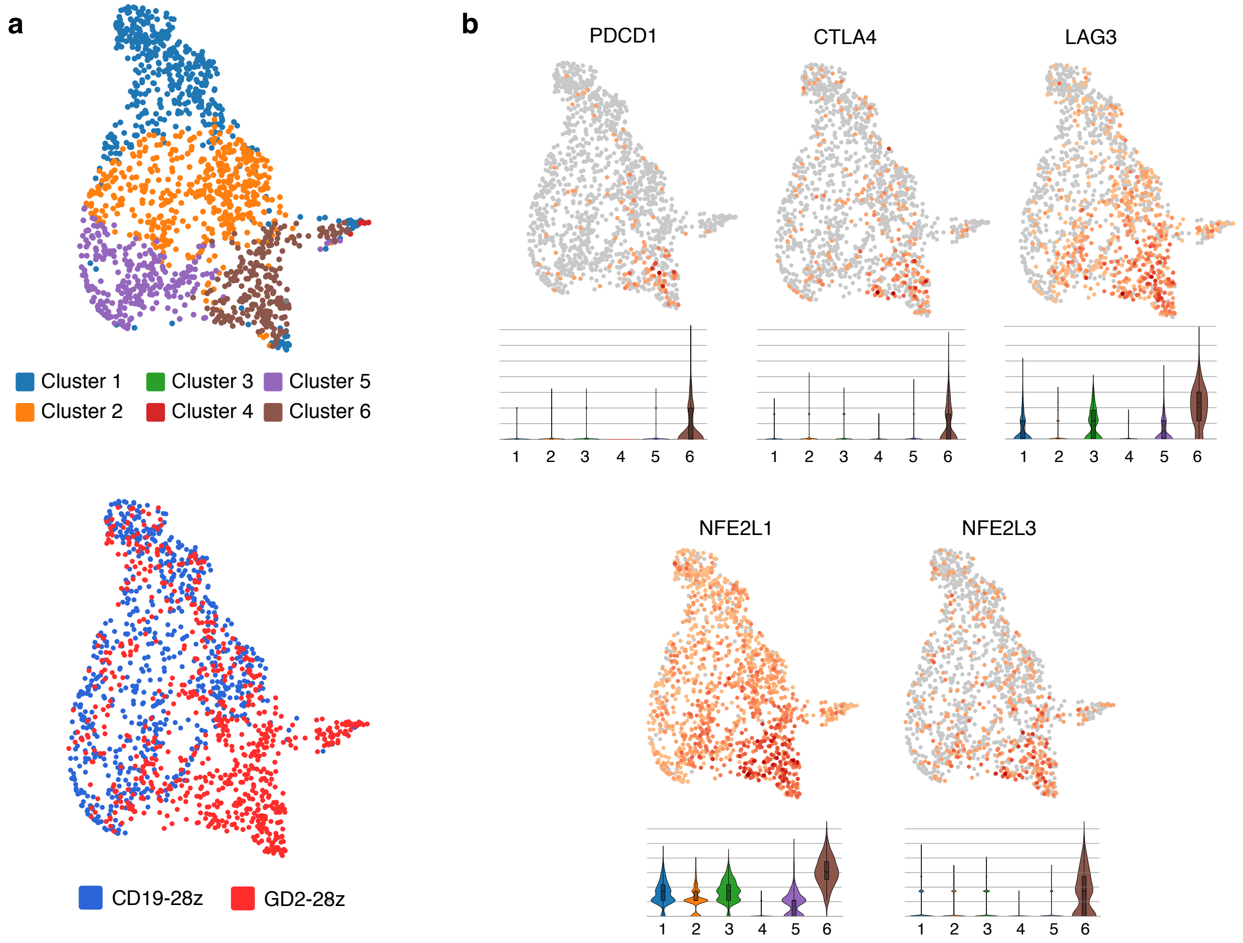


**Supplemental Figure 6. NFE2L1/NFE2L3 are co-expressed with markers of exhaustion.** (A) UMAP dimensionality reduction of single-cell RNA-seq on non-exhausted CD19-28z and exhausted HA-28z CAR T cells (from Lynn et al., 2019) identifies a cluster of cells unique to the exhausted CAR T cell population. (B) Inhibitory receptor markers of exhaustion (PD-1, CTLA4, LAG3) and NFE2L1/3 are expressed in the exhaustion-specific cell cluster.
